## Supplementary Materials for "MAIT cells contribute to protection against lethal influenza infection *in vivo*"

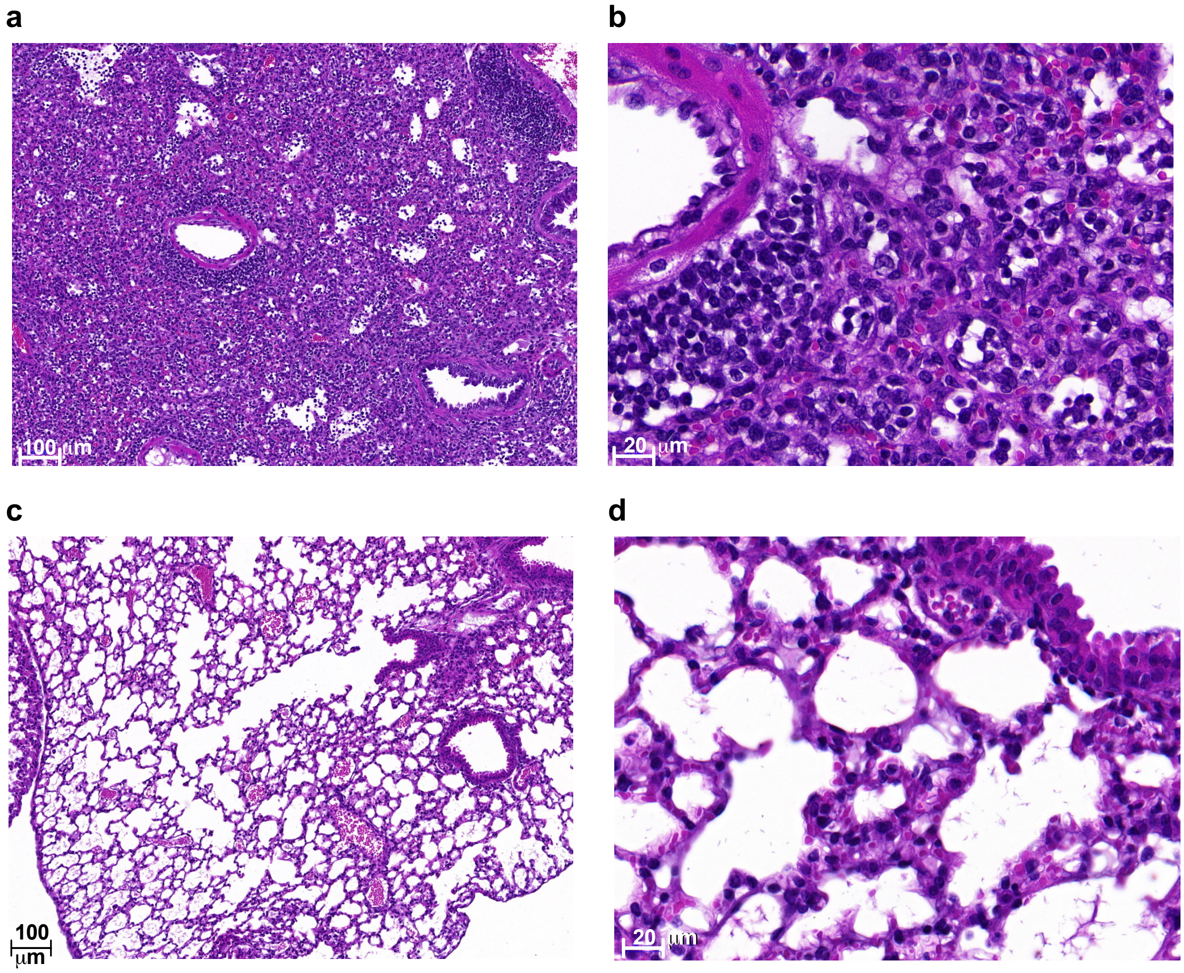


**Supplemental Figure 1. Pulmonary histology during influenza virus infection.**

Representative photomicrographs of haematoxylin and eosin-stained sections of lungs from C57BL/6 mice infected with 100 PFU of PR8 at 8 dpi. Images at low (**a**) and high (**b**) magnification show expansile parenchymal necrosis with severe perivascular, peribronchial and interstitial inflammation characterised by infiltrates of macrophages, lymphocytes and neutrophils. (**c, d**) Uninfected controls showing normal histology.

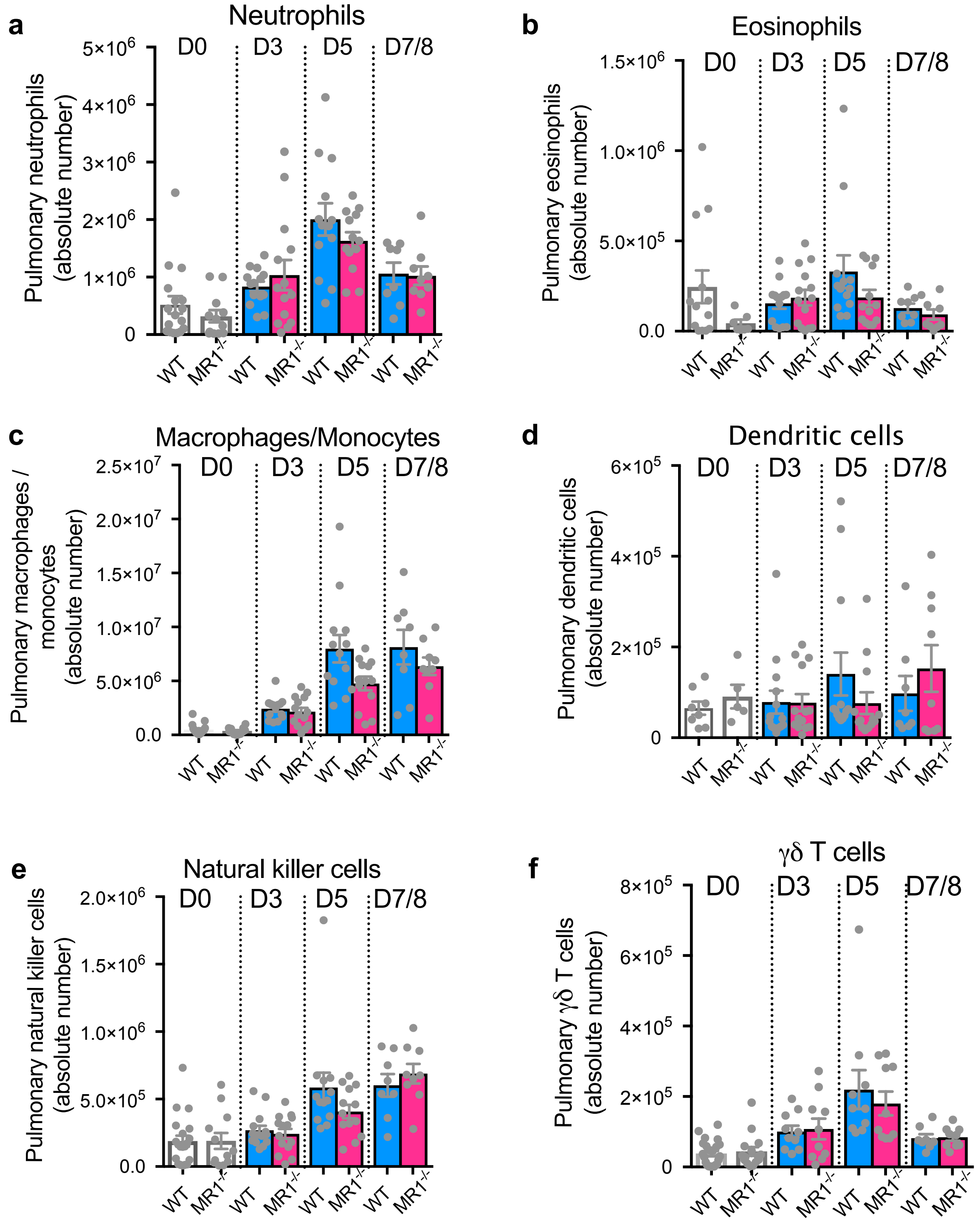


**Supplemental Figure 2. Innate immune cell recruitment in wild-type and MR1$^{-/-}$ mice during PR8 infection.**

(**a-f**) Absolute numbers of innate immune cells in the lungs of WT (C57BL/6) or MR1$^{-/-}$ mice before (D0) and at 3, 5 and 7 to 8 dpi (D3,D5,D7/8) following infection with 100-150 PFU of

PR8. (**a**) neutrophils (Ly6G$^{+}$SigF$^{-}$CD11b$^{+}$), (**b**) eosinophils (SigF$^{+}$CD11b$^{int}$CD64$^{-}$CD11c$^{-}$), (**c**) macrophages/monocytes (CD64$^{+}$CD11b$^{+/-}$Ly6C$^{+/-}$SigF$^{-}$), (**d**) dendritic cells (CD11c$^{+}$I-A$^{b+}$CD3$^{-}$CD64$^{-}$, (**e**) natural killer cells (NK1.1$^{+}$CD3$^{-}$CD19$^{-}$CD11b$^{-}$F4/80$^{-}$CD11c$^{-}$), (**f**) γδ T cells (TCRγδ$^{+}$CD3$^{+}$TCRβ$^{-}$CD19$^{-}$CD11b$^{-}$F4/80$^{-}$CD11c$^{-}$). Graphs shows means ±SEM. Combined data from two experiments with similar results, (total n=8-14 per group).

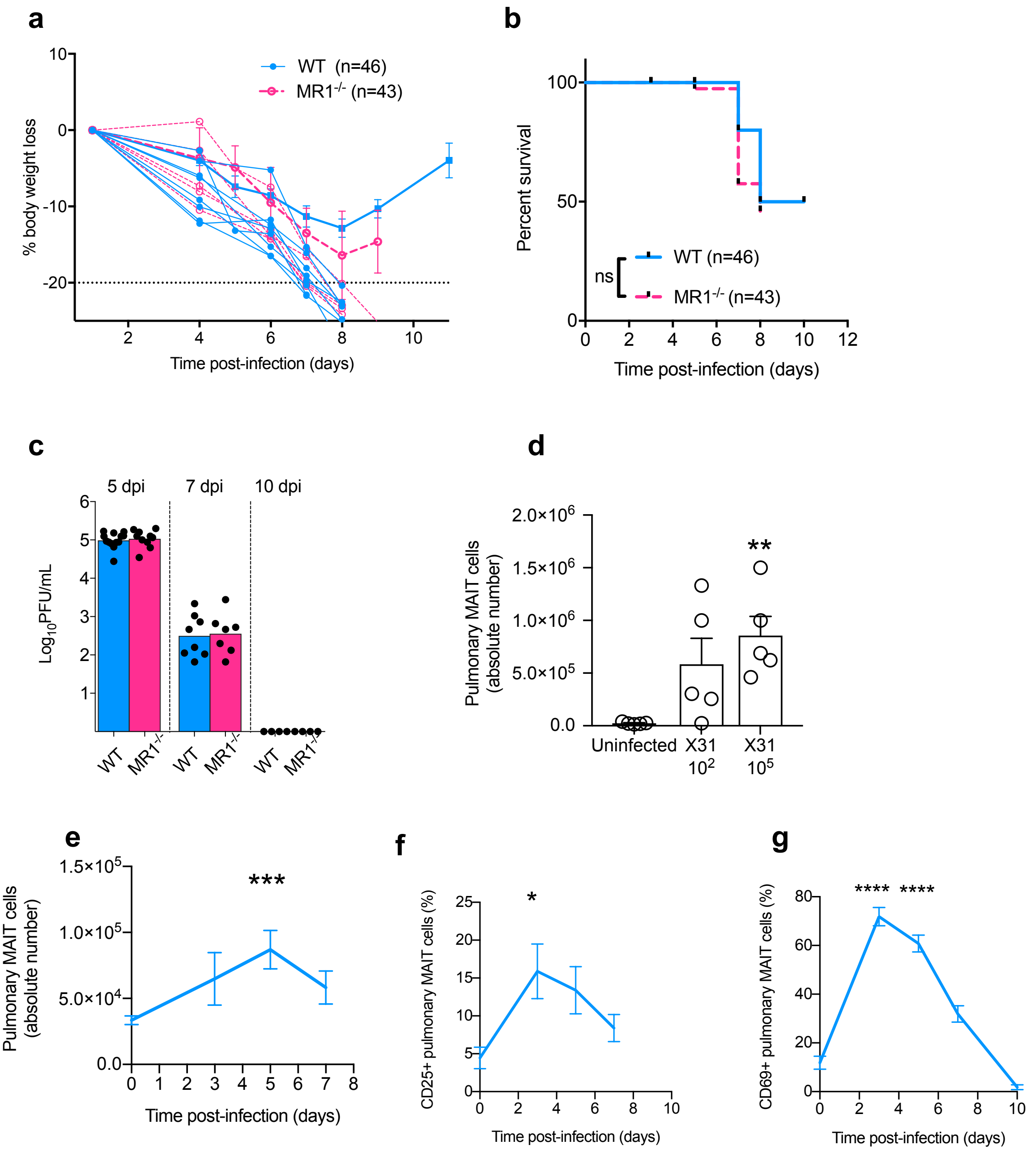


**Supplemental Figure 3. MAIT cell activation and accumulation after infection with the less pathogenic X-31 strain.**

(**a**) Body weight loss expressed as a percentage of starting weight and (**b**) survival curves after infection with 5000 PFU of X-31 virus. Data are combined from five experiments with similar results, comprising WT (n=46) and MR1$^{-/-}$ (n=43) mice. Weight loss graphs show mean weights±SEM for surviving mice, with individual plots for those which succumbed to infection. (**c**) Titres of infectious virus in clarified lung homogenates, expressed as PFU per lung at 5, 7 and 10 dpi. Data are from two independent experiments, each of n=3-6 / group. (**d**) Changes in absolute pulmonary MAIT cell numbers 7 days after intranasal infection with 100 or 10,000 PFU of X-31. Experiment performed once. (**e-g**) Absolute numbers of pulmonary MAIT cells (**e**) and proportions of MAIT cells expressing (**f**) CD25 and (**g**) CD69 in single-cell suspensions prepared from lungs before (D0) and after infection with 5000 PFU of X-31. Data are combined from three experiments with similar results. Graphs show means ±SEM. *, Mann-Whitney P<0.05.

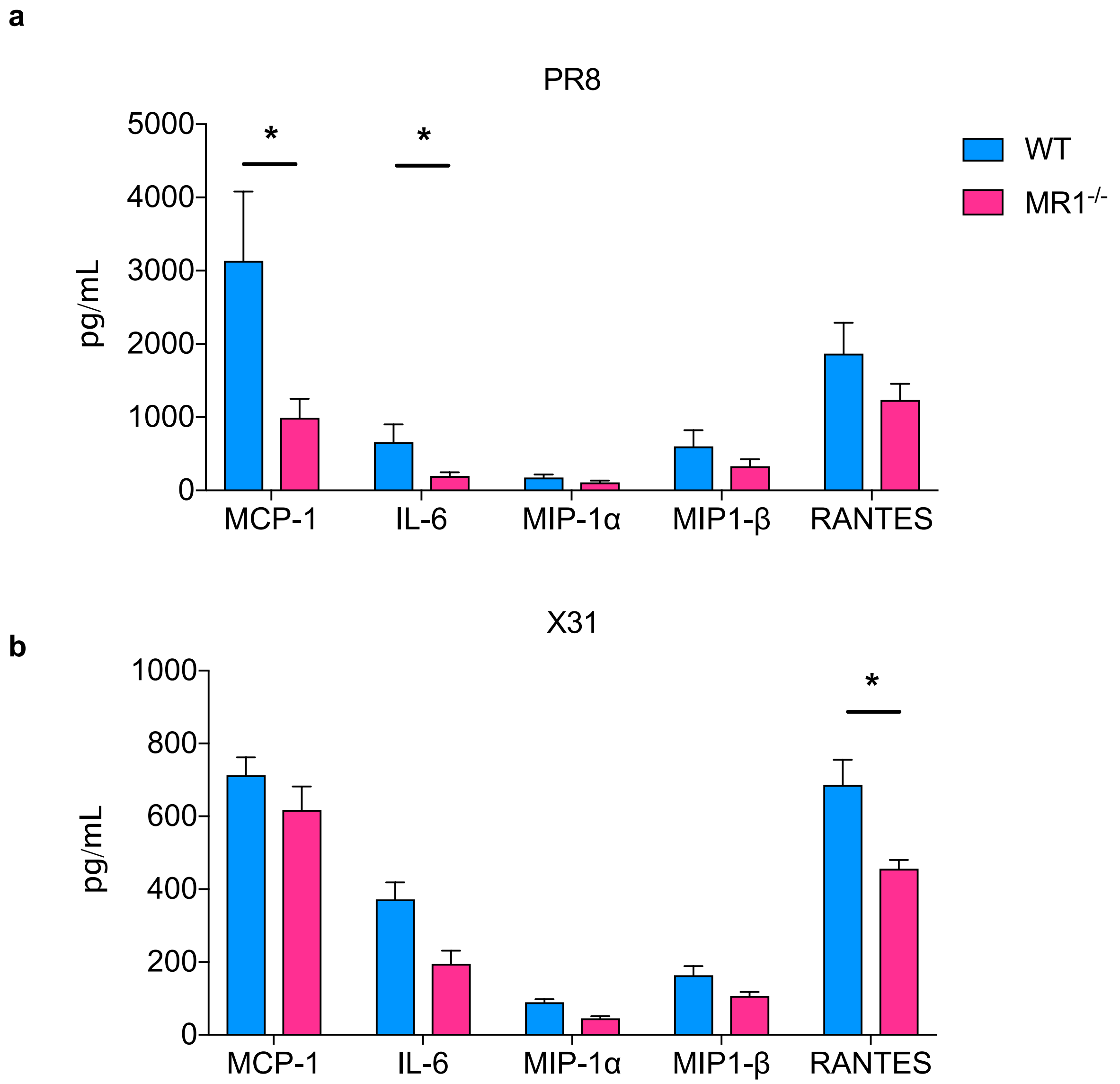


**Supplemental Figure 4. Innate inflammatory cytokines in wild-type and MR1$^{-/-}$ mice during influenza virus infection.**

Concentrations of key pro-inflammatory cytokines measured by cytokine bead array in homogenised lungs day 3 post-infection with (**a**) 100 PFU of PR8 and (**b**) 5000 PFU of X-31 compared in WT (C57BL/6) and MR1$^{-/-}$ mice. Mann-Whitney tests with Bonferroni correction for multiple comparisons. Graphs show means ±SEM of n=10 (PR8) or 5 (X-31) mice. *, $P<0.05$; **, $P<0.001$.
